# BlueNuclei: automated identification and classification of live and dead transfected neurons using interpretable features

**DOI:** 10.64898/2026.01.29.702657

**Authors:** Zhan Zha, Jing Jin, Russell L. Margolis, Daniel Taliun

## Abstract

*In vitro* modeling of neuronal disorders using transfected primary neurons is one of the fundamental approaches for studying disease mechanisms and therapeutic screening. Assessing neuronal viability is an everyday yet critical task in such experiments and requires accurate identification and classification of live and dead transfected neurons from dual-channel fluorescence images; however, this step is typically performed manually, resulting in inconsistent, labor-intensive, and poorly scalable analysis due to limitations of existing image analysis tools. Here, we present BlueNuclei, a user-friendly software with two modules: Hyades, which identifies nuclei of transfected neurons using dual-channel fluorescence image processing techniques, and Pleiades, an SVM-based classifier that distinguishes live from dead neurons using human-vision-inspired, biologically interpretable subnuclear features. Benchmarking on real images showed that BlueNuclei achieves near-human accuracy with substantially faster processing and minimal computational resources compared to deep learning alternatives when applied to the classification step. BlueNuclei provides a simple local user interface for data input and interactive visualizations that display classification results, including feature metrics and a confidence score for each nucleus. BlueNuclei offers the first scalable, fully automated, solution to viability assessment of transfected neurons, facilitating in vitro mechanistic studies of genetic neuronal disorders and therapeutic screening.

## Introduction

Many neurological disorders, such as Huntington’s disease, Charcot-Marie-Tooth disease, and spinocerebellar ataxias, are linked to genetic causes or risk factors^1-7^. Understanding how genetic mutations contribute to the pathogenesis of these diseases at the molecular level is a crucial step towards the discovery of drug targets and the development of therapies^3,8-11^. Robust *in vitro* model systems are essential for investigating how specific gene mutations affect neuron viability under both disease and healthy conditions ^12-18^. Primary neuron culture (PNC), typically derived from animal brains such as embryonic mouse brains, is one of the commonly used systems. PNC is favored for its relevance, efficiency, reproducibility, and versatility^14,16,17,19-21^, enabling the study of genetic effects on neuronal survival and pharmacological toxicity.

A neuronal toxicity assay includes *in vitro* neuron culture and transfection, image acquisition, and image analysis (**Figure 1**). Typically, primary neurons are isolated from embryonic mouse brains, transfected with a plasmid carrying the gene of interest and a fluorescent reporter gene (GFP for this study), and their nuclei are stained with a DNA-binding fluorescent dye, 4’,6-Diamidino-2-phenylindole (DAPI). Images are then acquired using a conventional fluorescent microscope, producing dual-channel images: the GFP channel highlights the cytoplasm of only successfully transfected neurons, while the DAPI channel shows the nuclei of all neurons. Image analysis follows in two consecutive stages. In the first stage (**Figure 1**, step 5), the territories of successfully transfected neurons are identified in the GFP channel and used to guide the identification of their corresponding nuclei in the DAPI channel. In the second stage (**Figure 1**, step 6), subnuclear features, such as intra-nuclear patterns, bright spots, and edge sharpness (**Table S1**), are used to classify these nuclei as either live or dead, allowing calculation of the overall death rate of transfected neurons. This paper focuses on image analysis (i.e. steps 5 and 6), with details of the PNC experiment and image acquisition provided in the **STAR Methods** section.

**Figure 1:**
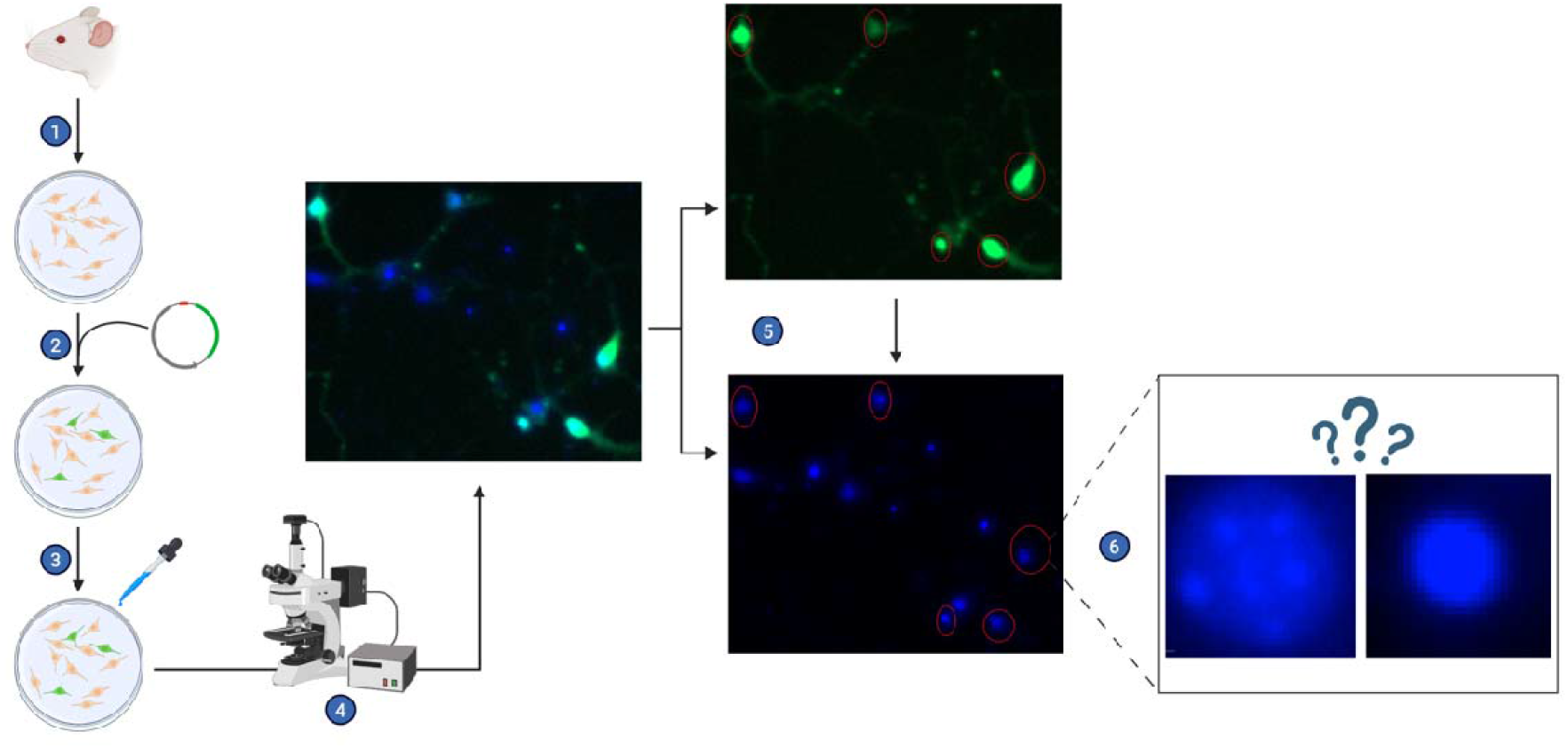
Overview of primary neuron culture (PNC) experiment and image analysis. Steps 1-4: experimental part (PNC and image acquisition); Steps 5-6: image analysis part. 1: Primary neurons are isolated from animal brains (e.g. mouse) and seeded onto a culture plate; 2: Neurons are transfected with a plasmid carrying GFP-tagged target gene. As a result, only successfully transfected neurons will express the GFP-tagged target and therefore emit green fluorescent signal; 3. All neurons are fixed by paraformaldehyde and stained by the nucleus fluorescent dye DAPI (regardless of whether successfully transfected or not, live or dead at the time); 4. Neurons are imaged using a conventional fluorescent microscope, producing a dual-channel (GFP and DAPI channels) image; 5. Nuclei of successfully transfected neurons are identified; 6. Each nucleus from step 5 is assessed and classified as either live or dead. In the example, the one on the left is live and the one on the right is dead. The death rate of transfected neurons can then be easily calculated. Figure created in https://BioRender.com.

Several popular Deep-Learning (DL)-based bioimage processing algorithms are partially relevant to either the first or second stage, but none is suitable as an all-in-one solution for the whole two-stage image analysis task in PNC. Among existing algorithms that are relevant to the first stage are the DL-based general-purpose cellular image segmentation tools Cellpose^22^, Cellpose-SAM^23^ and DeepCell^24^. These tools leverage multichannel image inputs (e.g., nuclear, cytoplasmic, or membrane signals) to improve segmentation accuracy in densely packed or overlapping cellular regions. In the context of the image analysis task in PNC, these methods can reliably segment all nuclei in the DAPI channel or all cellular territories in the GFP channel. However, they are not designed to identify and selectively associate individual nuclei in the DAPI channel with their corresponding cytoplasm territories from the GFP channel, nor to perform downstream viability classification on the selected nuclei. Thus, while these approaches are highly effective within their intended scope, they are not directly applicable to the specific cross-channel selection and nucleus-level classification required here without substantial task-specific re-engineering. The StarDist^25^ is another DL-based generalist model with general-purpose segmentation and classification functionalities for any star-convex objects. In the context of PNC, it can segment all nuclei in the DAPI channel and, in principle, assign each segmented nucleus to a “live” or “dead” class via a user-trained classifier. However, this classification step presumes the existence of a validated training model capable of distinguishing live from dead nuclei in transfected neurons, which is currently unavailable. Similar to Cellpose, Cellpose-SAM, and DeepCell, StarDist provides no mechanism for selectively identifying the subset of nuclei corresponding to cytoplasmic territories detected in the GFP channel.

Even if adopted to PNC image analyses, Cellpose/Cellpose-SAM, DeepCell, and StarDist typically assume a computer equipped with a dedicated GPU and accompanying software development tools (e.g. CUDA^26^, TensorFlow^27^, PyTorch^28^) for efficient processing of large (>5,000 × 5,000 pixels) images, which is not always accessible or easy to set up on standard PCs. Their results are inherently difficult to interpret due to the black-box nature of DL models^29,30^. Moreover, although Cellpose, Cellpose-SAM, and DeepCell offer publicly accessible web-based user interfaces, they are primarily intended for demonstration or exploratory use on small example images and are not designed for scalable analysis workflows, while StarDist doesn’t offer a user interface at all. Practical application of these methods to large datasets require familiarity with coding and software development environment installation and configuration, which limits their accessibility for experimental biologists with limited programming experience.

Here, we introduce BlueNuclei, a software tool with an intuitive user interface integrating two connected modules to perform both the first and second stages of the image analysis in neuronal toxicity assays in a single pass (see **STAR Methods**). The “Hyades” module identifies nuclei of transfected neurons using dual-channel images, while the “Pleiades” module is an SVM-based classifier trained on images from PNC experiments. “Pleiades” leverages five biologically interpretable subnuclear features—spottiness, spot distribution, edge gradient, area, and intensity—explicitly designed to capture the visual cues used by human experts to distinguish live from dead nuclei (**Table S1**). We compared Pleiades to a DL-based classifier implemented in StarDist, with both trained on the same dataset to ensure a fair comparison. Evaluation on unseen images showed that Pleiades produced near-human classification results, with lower error rates and significantly faster runtime than StarDist (see **Results-Prediction Accuracy**). Overall, BlueNuclei provides the first scalable, fully automated, all-in-one solution for neuron-specific nuclei identification with subsequent classification into live and dead using subnuclear features inspired by human visual cues, offering both accuracy and speed without requiring specialized hardware or programming experience.

## STAR * Methods

### The Hyades Module

As mentioned earlier, when given a dual-channel image (GFP and DAPI channels), the first thing (i.e. stage-1) BlueNuclei does is to identify the specific nuclei of successfully transfected neurons, which is different from a typical, general nuclei segmentation task. Therefore, we developed Hyades—a module that combines several optimized “classical” image processing techniques to do this task. In a nutshell, it first identifies the soma of transfected neurons in the GFP channel, then projects these soma territories to the DAPI channel and identifies the nuclear contours within them. For soma identification, existing neuronal morphology analysis tools introduce unnecessary computational overhead by focusing on neurite-specific features that are irrelevant to soma detection^31,32^. Therefore, instead of incorporating these tools, Hyades detects soma territories using the following process: It first implements threshold_triangle from the scikit-learn Python library^33^ to create a binary mask of the GFP channel. It then implements findContours from the OpenCV Python library^34^ on the binary mask to identify the contours of segmented “blobs” (i.e. neurons) and select them based on the following criteria: area > 400 pixels, circularity > 0.1. The process of thresholding and contouring finding is then repeated once, but this time by implementing threshold_otsu from the scikit-learn library and findContours with selecting criteria being: area > 200 pixels, circularity > 0.1. The identified neuronal contours from the two processes were then joined as a union. The pixel coordinates of each contour in this union are recorded and used for a geometric contour scaling:

Let 𝒰 be a neuron’s contour made up of *n* pixels:

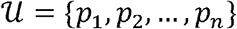

where ***p***_*i*_ = (*x*_*i*_,*y*_*i*_) ∈ *R*^𝟚^ represents the *i*^th^ pixels’ coordinates.

Let the centroid of the contour be:

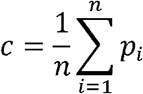

Let the scaling factor for the contour 𝒰 be:

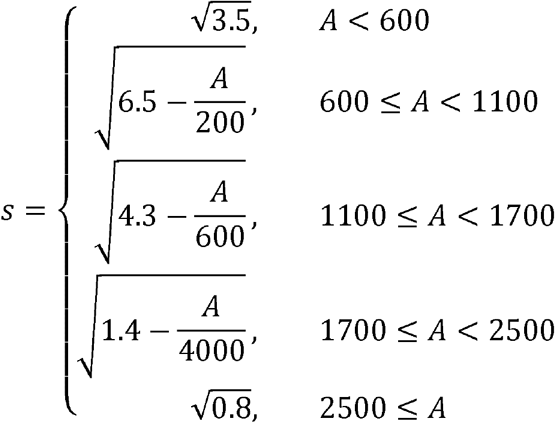

Where *A* is the enclosed area of 𝒰 (in pixels).

Then the new, scaled contour 𝒰 ^′^ for this neuron is:

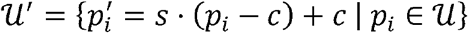

Where 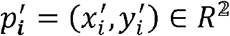 represents the *i*^th^ pixels’ coordinates and is integer rounded.

After finding and curating the neuronal contours, Hyades takes the DAPI channel image as input and implements GaussianBlur from the OpenCV Python library to blur the image with the following parameters: sigmaX = 27, sigmaY = 27, followed by the implementation of Laplacian from the OpenCV Python library to the blurred image with the following parameter: ksize = 3. Then it projects the neuronal contour pixels onto the DAPI Laplacian map and selects pixels from the DAPI channel that meet all the following criteria: 1. Intensity value > 2*M*, where *M* is the median greyscale intensity of the entire original DAPI channel; 2. Laplacian value > 0; 3. Residing inside neuronal contours. Among the selected pixels, small clusters of pixels (noise) are filtered out by implementing DBSCAN from the scikit-learn library with parameters: eps=2, min_samples=2.

After filtering, let ℰ be the candidate pixels within a neuronal contour 𝒰^′^:

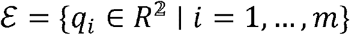

where *q*_*i*_ = (*x*_*i*_,*y*_*i*_) represents pixel *i*’s coordinates.

Let *c* be the centroid of the candidate pixels:

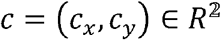

Then, group *q*_*i*_ in ℰ into 30 angular sectors around ***c***, and let each angular sector be:

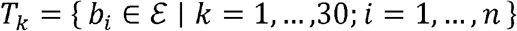

where *b*_*i*_ = (*x*_*i*_,*y*_*i*_) represents the coordinates of the *i*^th^ pixel within the *k*^th^ sector. Then *the* selected pixel from this sector is:

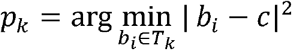

Let 𝒫 be a collection of those selected pixels:

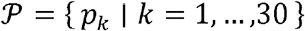

which represents the edge (i.e. contour) of the nucleus of neuron 𝒰^′^. The coordinates and intensities of pixels within the nucleus are extracted to calculate the five features of the nucleus (see **STAR Methods-Feature Calculations**). Eventually, each transfected neuron’s nucleus is transformed into a data frame with calculated features, which will be fed as input to the next part: nuclei classification (see below).

### The Pleiades Module

Pleiades is a linear SVM model trained (see **STAR Methods-Model Training)** to be able to use five human-knowledge-inspired features (see **STAR Methods-Feature Calculations** and **Table S1**) to classify the nuclei identified by Hyades as “live” or “dead” class. The classification is implemented simply using decision_function from the scikit-learn Python library. Note that Pleiades uses the same affine transformation formula and the same scalers during training and application to ensure consistent domain alignment of feature values of different numerical scales, which is critical for the program to work robustly on unseen images that might have different baseline conditions (e.g. brightness, resolution, etc.) than the training images.

## Feature Calculations

### 1. Spottiness

Spottiness refers to the abundance and prominence of subnuclear bright spots.

Let the entire image be a 2D Euclidean space of width *m* (x-axis) and height *n* (y-axis), where the upper-left pixel is at the origin (0,0). Each pixel has integer coordinates and an intensity value in the range [0,255].

Let a nucleus 𝒩 be a convex, filled polygon represented as a set of *N* pixels:

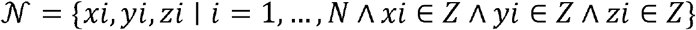

Where 0 ≤ *x*_*i*_ ≤ *m* and 0 ≤ *y*_*i*_ ≤ *n* are the corresponding 2D coordinates and 0 ≤ *z*_*i*_ *≤* 255 is the intensity value of a pixel *i*.

Let *f*_*x*_(*y*): ℤ → ℤ be a slice function defined for each 0 ≤ *x* ≤ *m* such that (*x,y,f*_*x*_(*y*)) ∈ 𝒩 . Then, for each 0 ≤ *x* ≤ *m*, we can define an ordered list of local maxima:

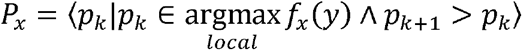

and an ordered list of local minima in between each pair of local maxima:

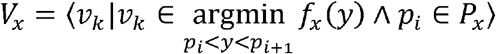

The spottiness score for each 0 ≤ *x* ≤ *m* can be defined as:

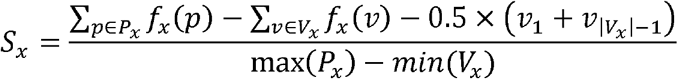

Analogously, we can define spottiness scores *S*_*y*_ for each 0 ≤ *y* ≤ *n*.

The final spottiness score for the nucleus is:

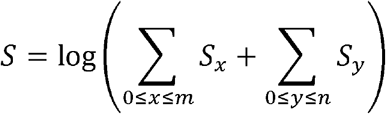

If *S*_*total*_ ≤ 0: then we assume that the nucleus has a very low degree of spottiness (i.e. very few internal bright spots) and therefore assigns *S* = −5 to avoid log error.

### 2. Spot distribution

Spot distribution is relevant to spottiness. It measures how spread out subnuclear bright spots are from the nucleus’s geometric center.

Let the set of 2D coordinates for all previously detected peak pixels in the nucleus 𝒩 be:

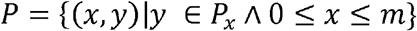

and let the centroid coordinates of the nucleus be:

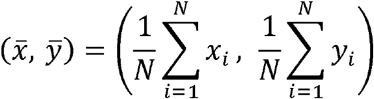

For each peak p ∈ *P*, compute its Eucledian distance to the centroid:

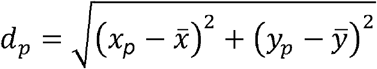

Final spot distribution score *D* is:

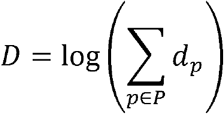

If:

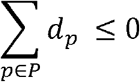

Then we assume that the nucleus has a very low degree of spot distribution of internal bright spots and therefore assigns *D* = −5.

### 3. Edge gradient

Edge gradient measures how “sharp” a nucleus’s edge is.

Let nucleus 𝒩 be 𝒩 = {(*x*_*i*_,*y*_*i*_) | *i* = 1,…,*n*} with centroid *C*. By implementing scale function from the Shapely^35^ Python library with the following parameters: xfact = 0.6, yfact = 0.6, we geometrically shrink 𝒩 to obtain a shrunken nucleus:

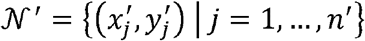

Let ℬ be the “belt” area that lies between the contours of the original nucleus and the shrunken nucleus, represented as a set of *B* pixels:

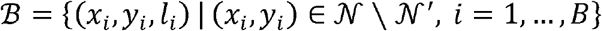

where *x*_*i*_,*y*_*i*_ are the coordinates, and *l*_*i*_ is the Laplacian value of the *i*^th^ pixel. Edge gradient (*E* ) is calculated as:

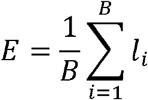

### 4. Intensity

Intensity refers to the average pixel value of the nucleus.

Let *M* be the median greyscale intensity of the entire input image, and, as mentioned above, let *N* be the total number of pixels within a nucleus, with the *i*^th^ pixel’s intensity value being *v*_*i*_. The intensity is calculated as:

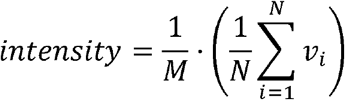

This measures the average brightness (normalized to the overall image brightness level) of the nucleus.

### 5. Area

Area refers to the total number of pixels within the nucleus. It is simply defined as the cardinality of set 𝒩:

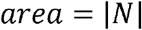

## Model Training

### PNC and transfection

To acquire images for training and testing the Pleiades model and the custom StarDist classification model, two PNC experiments, *E*1 and *E*2 were carried out by two different persons using the same experimental protocol. In each experiment, primary cortical neurons (with negligible presence of other cell types) from E16 CD1 mice were isolated and seeded on a 48-well plate with approximately 40,000-50,000 cells per well covered with 350μl of complete Neural Basal Medium with supplements (NBM), using a standard protocol^36^. Medium was fully replaced by fresh NBM at 24 hours and half-replaced at every other 48 hours post-isolation. Transfection was performed as previously described^37^. After transfection, cells were stained with DAPI (4′,6-diamidino-2-phenylindole, 0.2 μg/ml in 1X PBS) for 5 minutes protected from light at RT. Each well was washed with 500 μl of 1X PBS twice and covered in 500 μl of 1X PBS.

### Image acquisition

After transfection and staining, neurons were imaged using a Zeiss fluorescence microscope and the Zeiss Zen Pro software (v. 3.0) (Table 2). The focal plane position was manually set using the software for each well. Then the microscope’s built-in robotic camera automatically captured 36 small, rectangular, adjacent regions (i.e. “tiles”) from the central part of the well. The software then stitched the tiles into a single, large, dual-channel (DAPI and GFP channels) image and saved it as a CZI file (Zeiss microscope image file). As mentioned earlier, the DAPI channel is a grey-scale (pseudo-colored in blue) image that shows the DAPI-stained nuclei of *all* neurons (regardless of whether successfully transfected or not), while the GFP channel is a grey-scale (pseudo-colored in green) image that shows only the cytoplasm of *successfully transfected* neurons. The two channels were also saved as TIFF format for convenience. The CZI image files from *E*_1_ were named as *a*1, …, *a*8, *b*1, …, *b*8, *c*1, …, *c*8, *d*1, …, *d*8, *e*1, …,*e*8, *f*1, …, *f*8, while those from *E*_2_ were named as *t*_1_, …, *t*_48_. Because of low transfection efficiency, images from *E*_2_ were excluded from the *evaluation* of Pleiades’s and StarDist’s classification performance (see Methods-Performance Evaluation). Nonetheless, *E*_2_ images were pooled with *E*_1_ images for *training* both the Pleiades model and the custom StarDist model (see Methods-Model Training). Relevant metadata of *E*_1_ and *E*_2_ images can be found at **Table S2**.

### Training Pleiades

As mentioned earlier, Pleiades is a linear SVM model that classifies each nucleus as live or dead based solely on subnuclear features. For training, it only requires annotated nuclei from the DAPI channel, since the nuclear morphologies of transfected and non-transfected neurons do not differ systematically. In other words, the GFP channel plays no role in training Pleiades. Here we describe how we selected and annotated nuclei from the DAPI channels of *E*_1_ and *E*_2_ images to train Pleiades. Under this section, the term “image” refers to the DAPI channel, unless otherwise stated.

Given that images from the same experiment (either *E*_1_ or *E*_2_) tend to have the same resolution, dimensions, and exposure time, it is neither necessary nor feasible to label all nuclei from all images for training. Therefore, we arbitrarily picked 6 images from *E*_1_ and 5 images from *E*_2_ (total = 11 images), ensuring coverage of different experimental and optical conditions. For each image, we hand-picked a single crop-out region based on the following standards: 1. The region has roughly 20-40 nuclei in total; 2. The ratio of live/dead nuclei within the region is roughly 1:1 (**Figure S1**); 3. The region should not be visibly out-of-focus (i.e. blurry); 4. The region should contain both closely clustered nuclei and sparse nuclei. Each “crop” was then loaded into the QuPath software (0.5.1)^38^, where we hand-drew lines around each and every nucleus and manually labelled it as either “live” or “dead”, resulting in a total of 305 labelled nuclei **(Table S3)**. Out of these nuclei, 80% of them were randomly selected as the training set, while the rest 20% were used as initial model testing (data not shown). The coordinates and value of the pixels in each nucleus within the training set were extracted and used for computing the features (see **STAR Methods-Feature Calculations**). A data frame was then constructed, where each entry records a nucleus’s label (“live” or “dead”), origin (from *E*_1_ or *E*_2_), and calculated features. Because features of nuclei originated from each experiment forms distinct domain, we aligned the domains using affine transformation, with data from *E*_1_ set as the reference domain:

For each nucleus from *E*_2_, let *f* be the value of one of its calculated features:

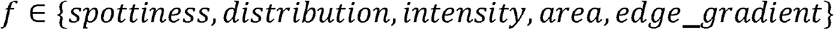

Then the feature’s value after domain alignment is:

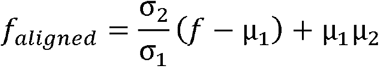

Where:

*σ*_1_, *σ*_2_ are the overall standard deviations of that feature in *E*_1_, *E*_2_, respectively

*μ*_1_,*μ*_2_ are the overall means of that feature in *E*_1_, *E*_2_, respectively

After performing domain alignment for *E*_2_ nuclei, we performed feature scaling for both *E*_1_ and *E*_2_ nuclei. Specifically, we applied the StandardScaler from scikit-learn Python library to normalize all features except the area feature, and the MinMaxScaler from the same library to the area feature and saved the two scalers as separate files. After feature scaling, the curated data frame was used to train Pleiades, a linear SVM model, by implementing the LinearSVC function from the scikit-learn Python library with the following parameters: C=5.0, max_iter=20000, dual=False, and class_weight=‘balanced’.

### Training StarDist

StarDist (v 0.9.1) is a general-purpose nuclei segmentation + classification tool based on deep neural networks (DNNs). It provides several pre-trained models for nuclei segmentation but lacks the ability to identify nuclei specifically belonging to transfected neurons (i.e. cross-channel referencing), nor does it include any model specifically for live–dead classification. To enable a fair comparison with BlueNuclei, we used the exact same 305 nuclei (**Table S3**) that were used for training Pleiades to train a custom StarDist classification model by following StarDist’s official guidelines and default settings.

Because StarDist cannot perform neuron-specific nucleus identification, we built a hybrid model combining our Hyades module (for identifying nuclei of transfected neurons) with the custom StarDist classification model. This hybrid (Hyades + StarDist) model was then benchmarked against our full BlueNuclei program (Hyades + Pleiades) (see **Results**) on the same test data (see **STAR Method-Test Data**).

### Test data

As mentioned earlier, experiment *E*2 had very low transfection efficiency, resulting in very few neurons in the GFP channel. Since identifying nuclei of transfected neurons is the first stage of the image analysis process, *E*2 images, although used in training the Pleiades and the StarDist nuclei classification models, are unsuitable for the purpose of testing. Therefore, the test data consists of 18 images arbitrarily picked from *E*1 only (a1, a2, a3, b1, b2, b3, c1, c2, c3, d1, d2, d3, e1, e2, e3, f1, f2, f3,) for the evaluation. We evaluated the results generated from BlueNuclei (Hyades + Pleiades) and StarDist (Hyades + custom StarDist classification model) in the classification task (i.e. stage-2) using human raters’ results as the gold standard (see **Results**).

## Results

### The BlueNuclei Program

The BlueNuclei program integrates two consecutive modules, the Hyades and the Pleiades modules, into a completely automated tool with a user-friendly web-based interface (**Figure 2**; **Figure S2**).

**Figure 2:**
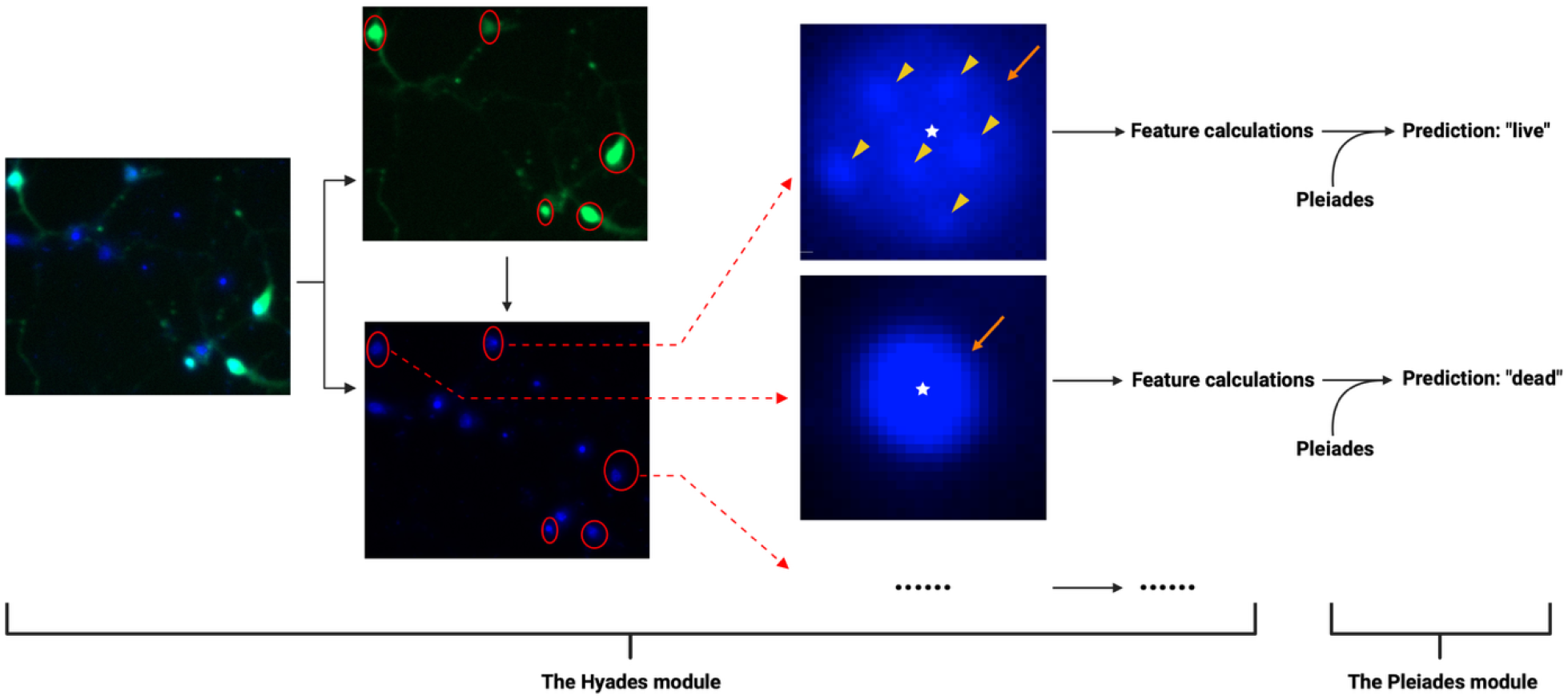
The BlueNuclei program (Hyades + Pleiades modules). First, Hyades takes a two-channel image, detects nuclei of transfected neurons (red circles) via cross-channel referencing, and extracts five features— spottiness (triangles), spot distribution (stars), area, intensity, and edge gradient (arrows). Then, these features are fed into Pleiades, a linear SVM model, to classify the nucleus as either dead or live (see STAR Methods for details).

The Hyades module takes a dual-channel image as input and selectively identifies nuclei of transfected neurons by first detecting cytoplasmic territories in the GFP channel using thresholding and shape-based filtering. During thresholding, it detects cytoplasmic contours across a range of GFP signal strengths by integrating Otsu’s method^39^ and triangle algorithm^40,41^. During shape-based filtering, it prunes out neurites and adjusts the sizes of cytoplasmic contours using morphological operations. Then, the identified cytoplasmic contours are projected onto the DAPI channel to delineate their enclosed nuclear contours based on high-contrast edge pixels. (see **STAR Methods-The Hyades Module**). The coordinates of these nuclei will be ported to the Pleiades module.

The Pleiades module predicts whether a nucleus is “live” or “dead” using a supervised linear SVM classifier trained on five subnuclear features designed to mimic human visual assessment: spottiness, spot distribution, edge gradient, area, and intensity (see STAR Methods—The Pleiades Module). All five features were designed to quantitatively emulate visual cues used by human annotators when assessing nuclear viability. While area and intensity correspond to directly measurable image properties, the remaining three features—spottiness, spot distribution, and edge gradient—were defined in this study, with explicit quantitative formulations, to capture the corresponding qualitative visual patterns used by human observers. After expressing all five features as quantitative metrics, we evaluated their discriminative power for distinguishing live from dead nuclei. Specifically, using the annotated nuclei included in model training (n = 305), we individually assessed their distributions in live and dead nuclei after domain alignments (**Figure S3**). All five features significantly differentiated live and dead nuclei (two-sided Wilcoxon rank-sum test; P-value ≪ 0.0001), with large effect sizes (Cohen’s d between 1.80 and 4.64), indicating strong differentiation between the two classes (i.e. live vs dead).

### Prediction Accuracy

Since no existing freely available tool other than BlueNuclei is suited to the full two-stage analysis, we focused on benchmarking only the classification component. For a fair comparison, we evaluated the classification results (i.e. neuronal death rates) of BlueNuclei (Hyades + Pleiades) with a custom StarDist-based workflow (Hyades + a StarDist classification model we trained using the same training data as Pleiades) (**Figure 4**). For each of the 18 test images, BlueNuclei, StarDist, and three human annotators independently estimated the neuronal cell death rate, with the mean of the human estimates treated as the reference. We quantified each algorithm’s deviation from the human reference (i.e. classification error) as the absolute arithmetic difference in estimated cell death rate. Deviations across all images were pooled to compute the mean absolute error (MAE) of each algorithm.

We found that BlueNuclei has significantly lower classification error (MAE = 0.0400±0.0060 (SEM)) compared with StarDist (MAE=0.0985±0.0184(SEM)), respectively; Paired Wilcoxon signed-rank test; P-value = 8.39x10^-4^) (**Figure 3 and Table S4**).

**Figure 3:**
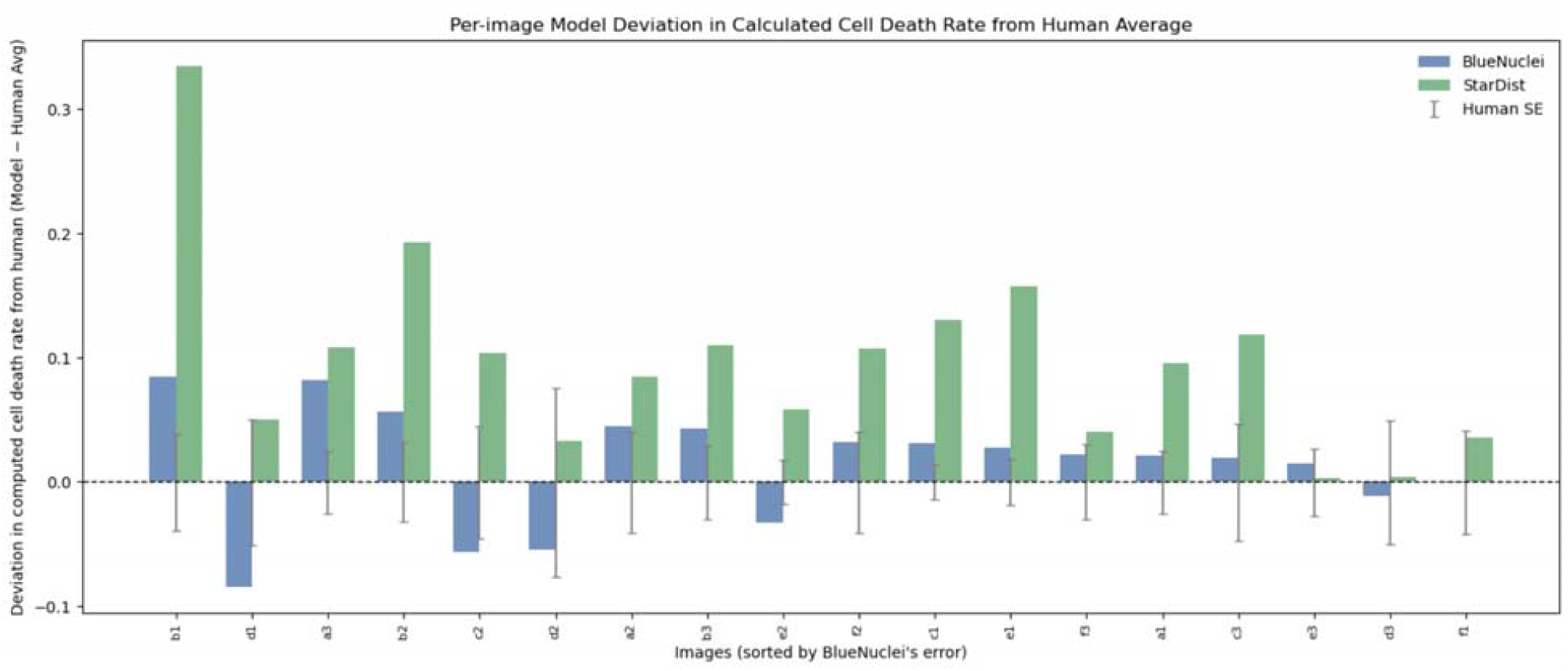
Per-image deviation of BlueNuclei’s vs StarDist’s cell-death rate estimates relative to human raters. Predicted cell-death rates for transfected neurons were computed by BlueNuclei (blue) and StarDist (green) across 18 test images and compared to three independent human raters. For each image (x-axis), the human-averaged cell-death rate is treated as the reference (dashed horizontal line at y = 0). Bar heights indicate the signed deviation of each algorithm’s estimate from the human average (algorithm − human average). Vertical gray error bars represent the standard error of mean (SEM) across the three human raters, reflecting inter-rater variability for each image.

We also found that in 14 of the 18 images (78%), BlueNuclei’s deviation from the human estimate was smaller than that of StarDist, whereas the converse was observed in only 4 images (22%), a difference that is statistically significant (binomial test against a null hypothesis H_0_:p = 0.5; P-value = 0.03) (**Table S4**).

We further evaluated each algorithm by counting the number of images in which its estimated cell death rate fell within one SEM of the human estimate. BlueNuclei achieved this in 8 of 18 images (44%), whereas StarDist did so in 5 of 18 images (27%) (two-tailed binomial test, P-value = 0.09; **Table S4**).

Overall, these results highlight the lower classification error and greater consistency with human raters of BlueNuclei’s classification performance on the test images.

### Speed

We next compared the speed of BlueNuclei and StarDist on various machines using 18 images, each with a resolution of 6,688 x 5,654 pixels and pixel size of 0.345 x 0.345 μ m. On average, BlueNuclei ran 1,000 to 14,000 times faster during model training phase (**Table 1A**) and 4 to 360 times faster during application (**Table 1B**) than StarDist, depending on the operating machine.

**Table 1:**
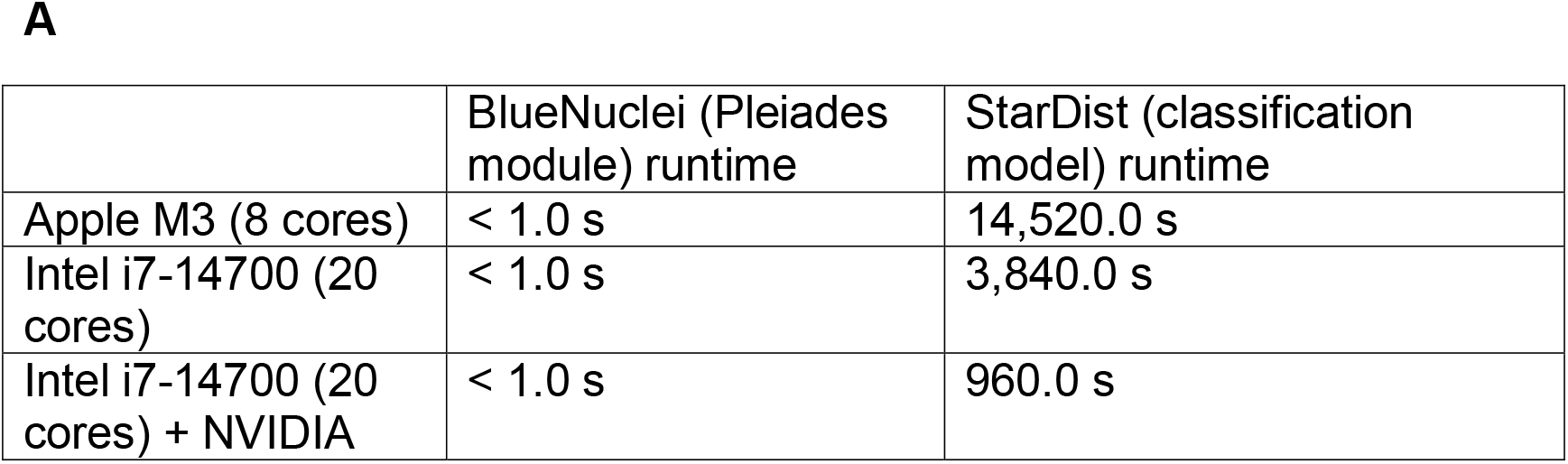
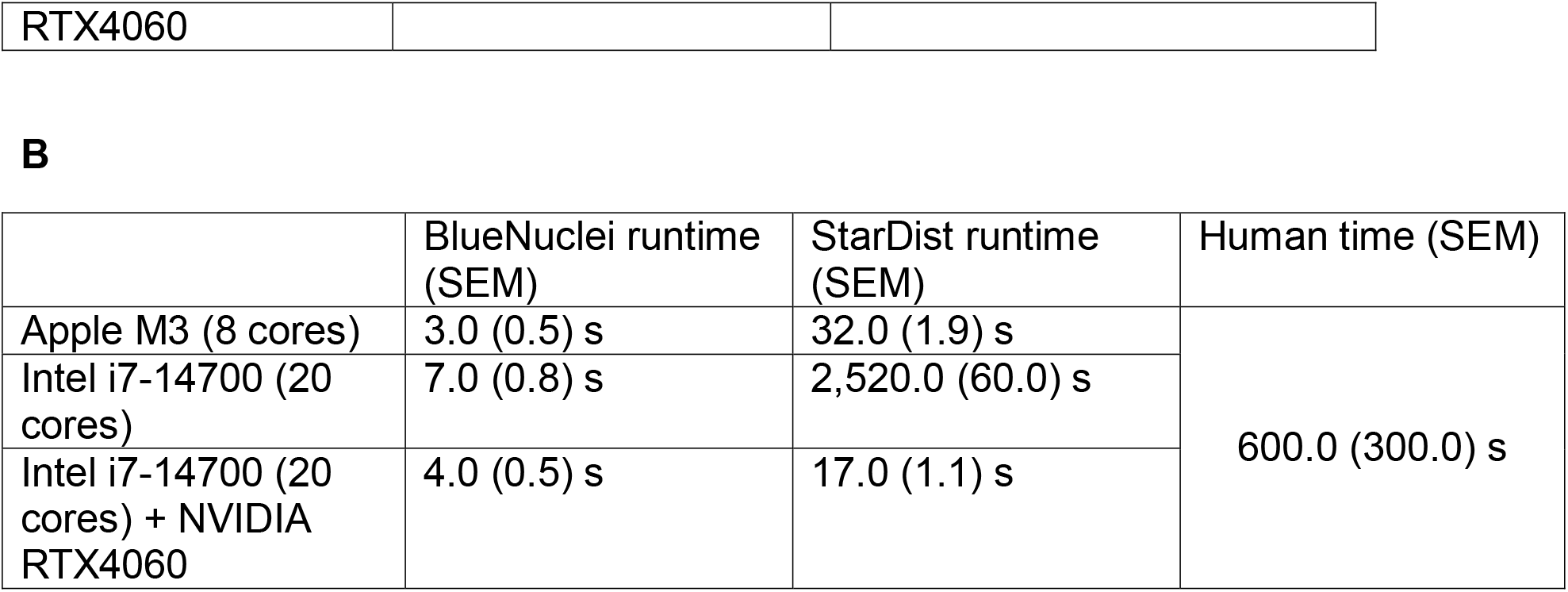
Comparing the speed of BlueNuclei vs StarDist during training and application on different machines. **A:** Time consumed to train BlueNuclei’s Pleiades linear SVM model and StarDist’s DL-based classification model during the training phase (305 nuclei in total). **B:** Average application run time (without visualization) per image for the BlueNuclei program (Hyades + Pleiades), the custom StarDist workflow (Hyades + StarDist classification model), and human. The algorithms’ run time is expressed as average across 18 testing images, while the human time is an estimation based on practical experience from the three human raters.

## Discussion

BlueNuclei is a tool capable of identifying and classifying live and dead transfected neurons in dual-channel fluorescence microscopy images from primary neuron culture experiments (**Figure 2**). Identifying transfected neurons poses unique challenges: (1) neurons exhibit diverse morphologies and irregular shapes; (2) cultures are typically dense, leading to frequent overlap among cells; (3) extensive neuronal processes (axons and dendrites) often traverse multiple nuclei in the DAPI channel and can be mistakenly recognized as soma; and (4) transfected neurons show highly variable GFP signal intensities. The Hyades module of BlueNuclei addresses these challenges by using double-thresholding technique combined and morphological refinements to increase the sensitivity and precision in capturing the soma of neurons with weak GFP signals without over-detecting those with strong GFP signals, at the same time of minimizing interfering signals from neurites or nearby neurons in cell-dense regions (see STAR Methods).

For classification, BlueNuclei’s Pleiades module is a supervised SVM model trained to use human-vision-inspired subnuclear cues to distinguish dead from live neurons. In the tested dataset, Pleiades shows lower error rates, faster speed and greater consistency with human raters than the tested alternative DL-based algorithm that could be adopted for the same purpose (**Figure 3, Table 1**, and **Table S4**).

BlueNuclei’s locally hosted web-based user interface (UI) allows researchers to quickly provide images for the analyses, produce both visual and quantitative and visual identification and classification results within a few seconds, and interactively explore them (**Figure S2**). In the UI, each predicted nucleus and its corresponding cytoplasm are highlighted and annotated with feature values and a classification confidence score. Borderline classification cases (such as a “dying” neuron) with low confidence scores are further highlighted, allowing users to manually inspect them and understand which feature(s) set them apart from the typical death and live classes.

This work has several limitations. First, BlueNuclei has been evaluated using only 18 images from two experiments, both with relatively narrow ranges of ground-truth cell death rates; thus, further testing on larger and more diverse datasets is required to assess the generalizability of the results. Second, only a few transgene constructs were used in the experiments, so it is unclear whether the five subnuclear features used by BlueNuclei’s SVM will retain similar discriminative power when applied under other experimental conditions. Third, BlueNuclei is specifically optimized for neuronal morphology, its direct applicability to other cell types may be limited without additional retraining or feature adaptation.

Despite these limitations, BlueNuclei is a useful tool that enables primary neuron image analysis at scale by shortening the whole process to a few clicks, avoiding the otherwise time-consuming manual task. Future expansion to larger datasets, additional experimental conditions, and other file formats may further broaden its applicability beyond neurons. As a proof of concept, BlueNuclei demonstrates how biologically grounded subnuclear feature design and lightweight machine learning can enable accurate, scalable, and interpretable image-based analysis in neurobiology.

## Abbreviations

PNC: primary neuron culture
DL: Deep Learning
SVM: Support Vector Machine
GFP: green fluorescent protein
DAPI: 4′,6-Diamidino-2-phenylindole

## Software Availability

The BlueNuclei software together with its source codes and models, the StarDist-based live/dead classification model, a Python notebook about deploying the custom StarDist workflow (Hyades + StarDist classification), and a Python notebook about training a custom Pleiades module are all freely available at: https://github.com/ZHAN-ZHA/BlueNuclei.

## Data Availability

A small part of the data is provided together with the code as an example. The complete training and testing data are available upon request.

## Acknowledgements

We thank Dr. Scot Kuo (Director of Microscope Facility, Johns Hopkins University) and Dr. Kwame Kutten for consultation on image analysis and Dr. Hanna Jaaro-Peled and Dr. Steven Doll (Laboratory of Genetic Neurobiology, Johns Hopkins University) for useful discussions. We appreciate GitHub suggestions from Sedat Demiriz and Dr. Vincent Chapdelaine (Daniel Taliun lab, McGill University).

