## Supplementary Information for "BlueNuclei: automated identification and classification of live and dead transfected neurons using interpretable features"

| **Feature** | **Description** | **Live cell** | **Dead cell** |
| --- | --- | --- | --- |
| Spottiness | The abundance and prominence of “bright spots” within the nucleus | Relatively high | Relatively low |
| Spot distribution | The average Euclidean distance between the “bright spots” and the nucleus’s centroid | Relatively high | Relatively low |
| Edge gradient | The sharpness of the nucleus’s edge | Relatively low | Relatively high |
| Area | Normalized size of the nucleus | Relatively big | Relatively small |
| Intensity | Normalized average brightness of the nucleus | Relatively low | Relatively high |

**Table S1. Distinguishing morphological features of live and dead nuclei of neurons.** These features, inspired by human domain knowledge, were used to train the Pleiades module to distinguish live and dead neurons (see STAR Methods for details).

| Experiments | Total number of images | Bit depth | Magnification | Aperture | Width of each image(px) | Height of each image (px) | Resolution (μm/px) | Exposure: DAPI (ms) | Exposure: GFP (ms) | Transfection efficiency |
| --- | --- | --- | --- | --- | --- | --- | --- | --- | --- | --- |
| E1 | 48 | 16-bit | 20 | 0.4 | 6688 | 5654 | 0.345 | 180 | 400 | ~5% |
| E2 | 48 | 16-bit | 40 | 0.65 | 13552 | 11308 | 0.172 | 316 | 521 | ~1% |

**Table S2. Images from experiments E1 and E2.** These images were utilized in model training and evaluation (see Methods-Model Training for details)

| Crop from image (experiment) | Number of labelled live nuclei | Number of labelled dead nuclei | Number of total labelled nuclei |
| --- | --- | --- | --- |
| *a1 (E1)* | 18 | 18 | 36 |
| *b1 (E1)* | 10 | 7 | 17 |
| *c1(E1)* | 16 | 20 | 36 |
| *d1(E1)* | 17 | 12 | 29 |
| *e1(E1)* | 15 | 19 | 34 |
| *f1 (E1)* | 13 | 17 | 30 |
| *t1 (E2)* | 8 | 16 | 24 |
| *t5 (E2)* | 14 | 10 | 24 |
| *t10 (E2)* | 10 | 13 | 23 |
| *t15 (E2)* | 10 | 8 | 18 |
| *t20 (E2)* | 12 | 22 | 34 |
| *Total (E1)* | 89 | 93 | 182 |
| *Total (E2)* | 54 | 69 | 123 |
| *Total (All)* | 143 | 162 | 305 |

**Table S3. Image crops used for model training.**

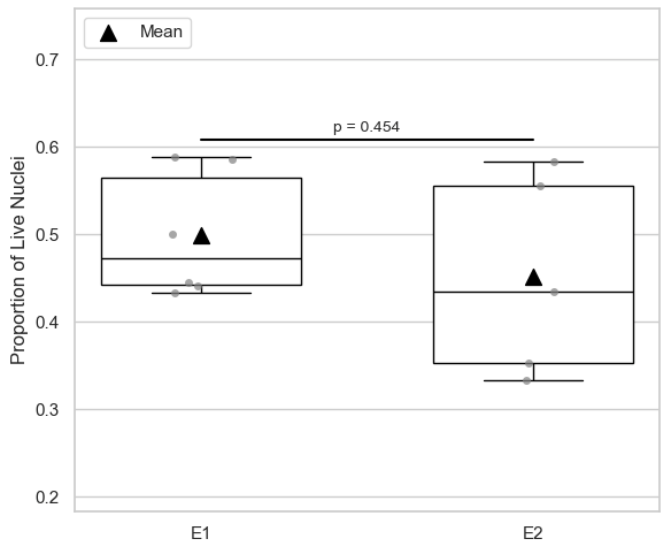

**Figure S1. Proportions of live and dead nuclei from E1 and E2 training crops.** Top/bottom whiskers: maximum and minimum. In-box middle line: median. Triangle: mean. The p value is computed using t-test of means. An insignificant p-value (p=0.454) indicates that live and dead nuclei are equally represented in both E1 and E2 crops.

**A**

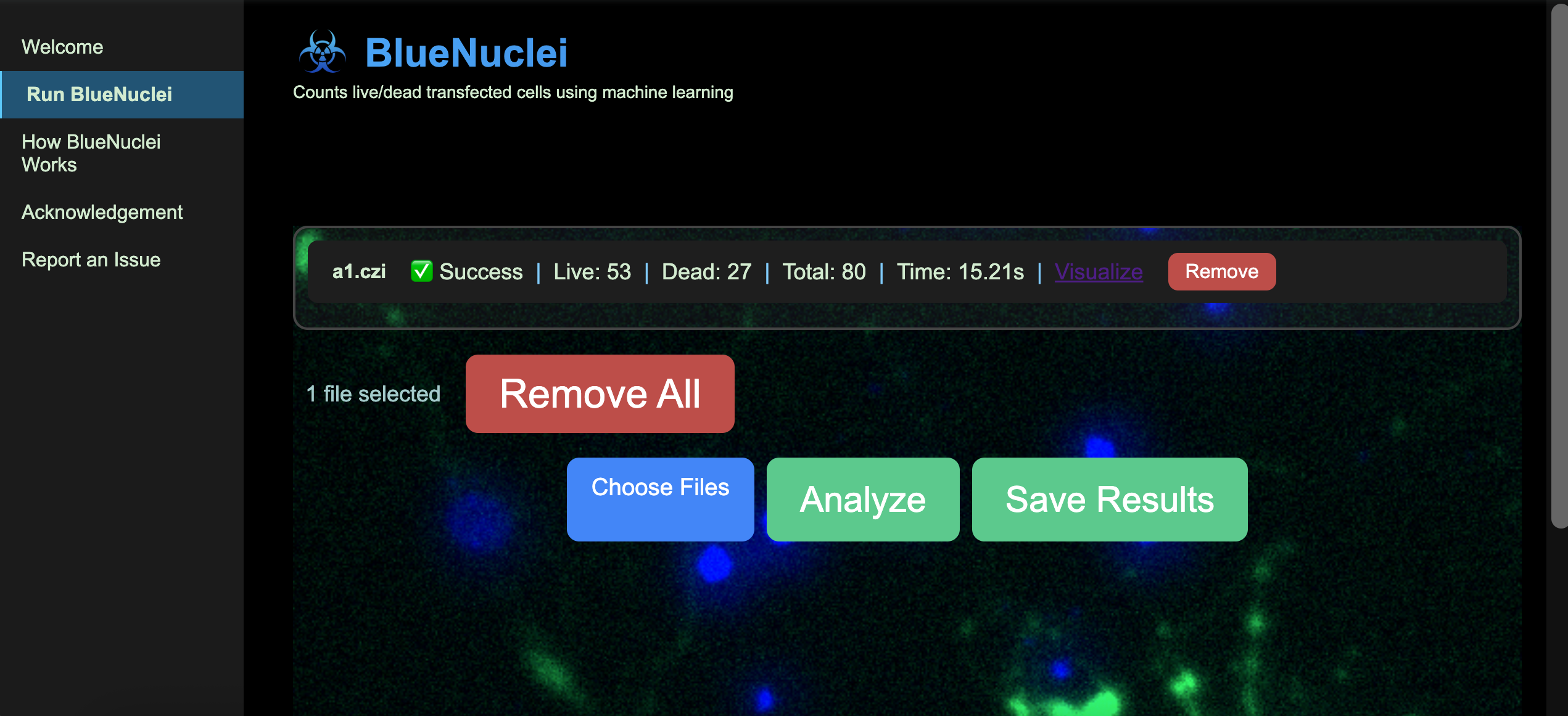

**B**

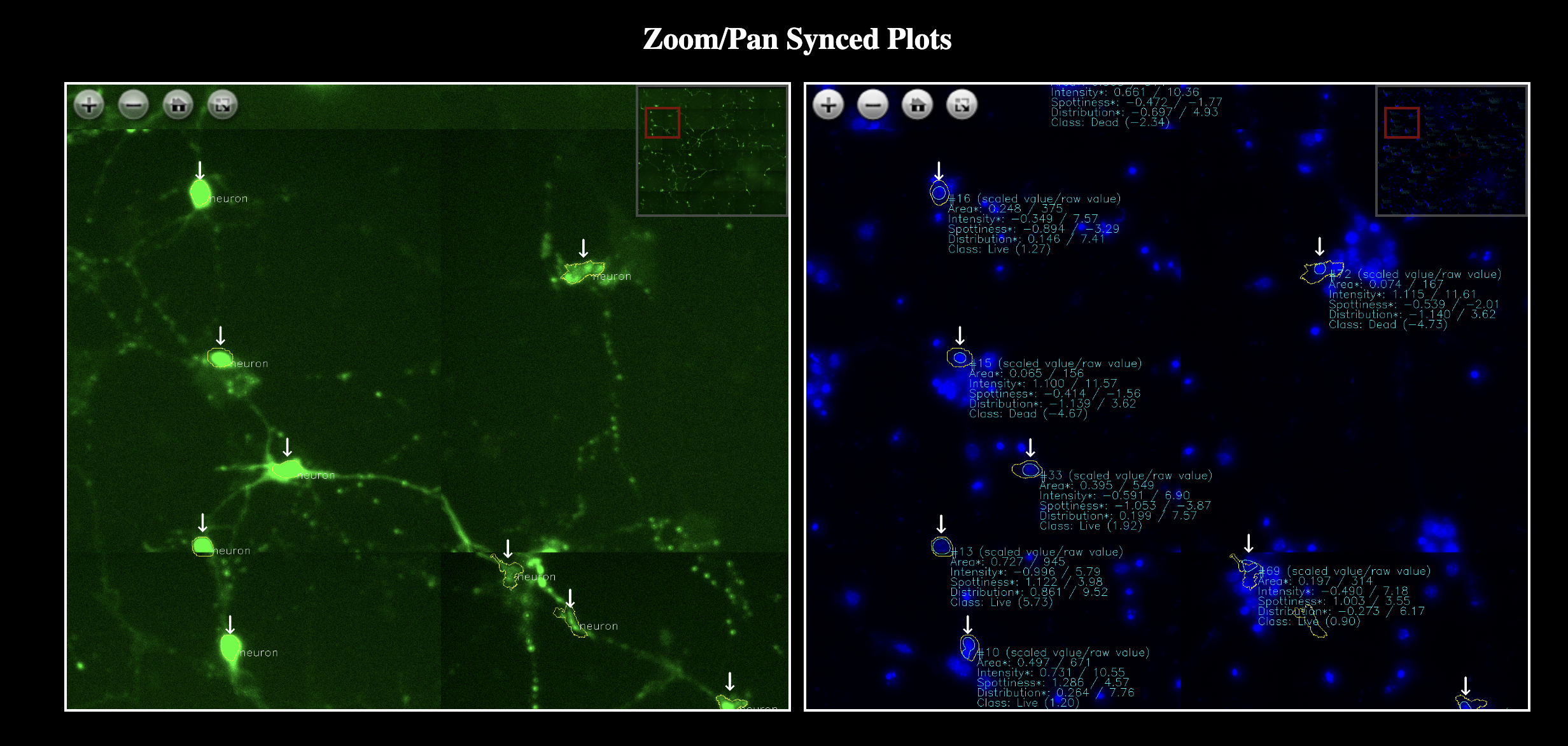
 **Figure S2. The user interface of BlueNuclei program. A:** The main interface where users upload images, run analysis and obtain results (i.e. count of live/dead transfected neurons). **B:** The visualization interface where transfected neurons identified by the Hyades module (GFP channel, left panel) and their corresponding nuclei (DAPI channel, right panel) are synchronously shown. The calculated feature values are also displayed for the nuclei in the right panel. Users can zoom in the two panels synchronously to see details of each predicted instance.

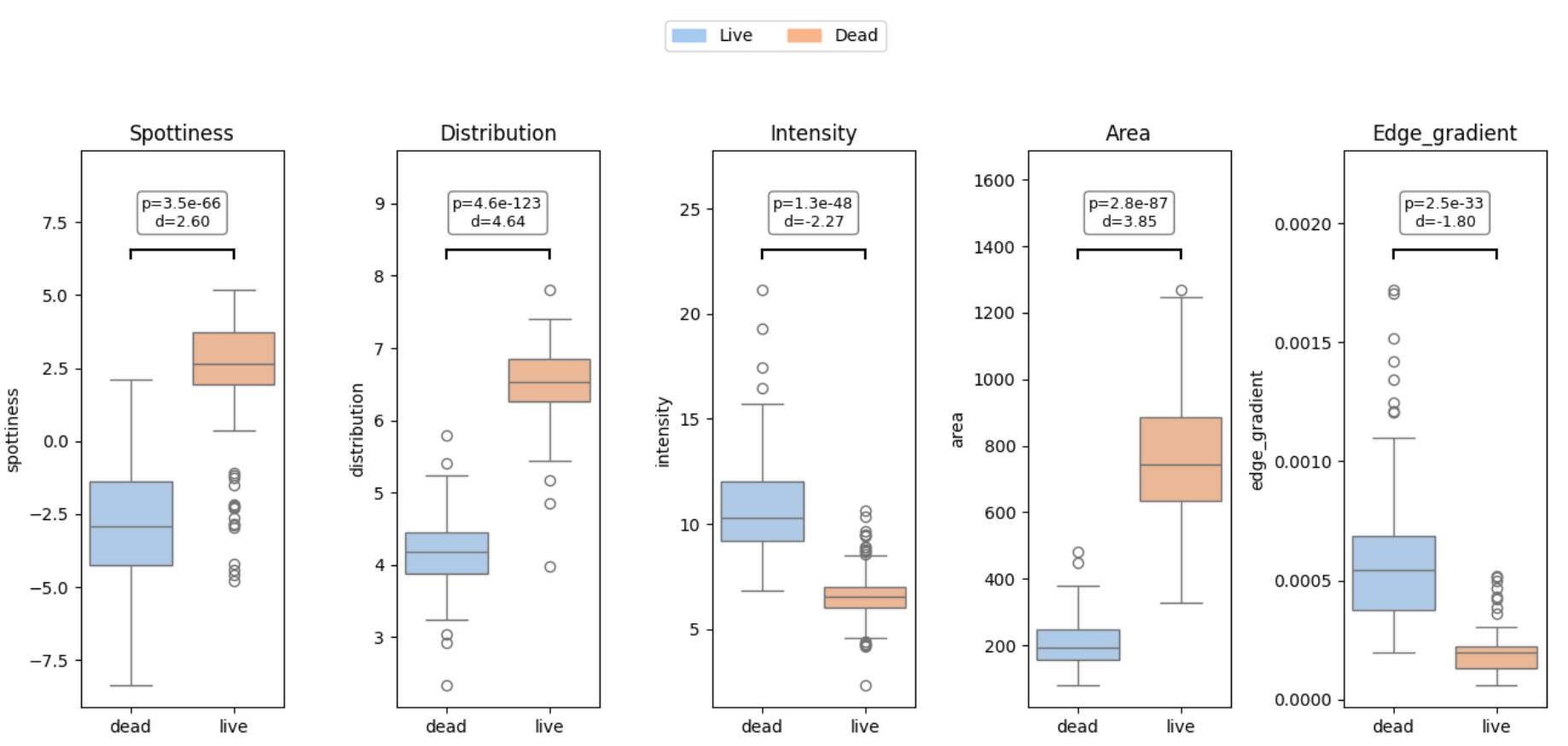

**Figure S3. Domain-aligned features of live and dead nuclei in the training data.** The five features (spottiness, spot distribution, intensity, area, and edge gradient) are computed from each nucleus from the training data (305 nuclei in total). Since nuclei from different image crops might have different optical conditions and thus differ in numerical scales, domain alignment was performed for each feature using affine transform (see STAR Methods for details). The distribution of these features are shown here to demonstrate their power at at discriminating live (blue) from dead (orange) nuclei. The feature values are displayed here in original scales. Circles: outliers. Top/bottom whiskers: maximum and minimum. In-box middle line: median. The p values (p) and Cohen’s d values (d) are displayed.

|  | BlueNuclei | StarDist | P value | Test Type |
| --- | --- | --- | --- | --- |
| MAE (±SE) | 0.0400 (±0.0060) | 0.0985 (±0.0184) | 8.39x10^-4^ *** | Wilcoxon |
| Closer to Human | 14 | 4 | 0.03 * | Binomial (two-tailed) |
| Within Human SE | 8 | 5 | 0.09 | Binomial (one-tailed) |

**Table S4. Comparison of BlueNuclei and StarDist error rates in neuronal cell death estimation.**
For each of the 18 test images, BlueNuclei, StarDist, and three human raters independently estimated the cell death rate of transfected neurons. The mean of the three human estimates was treated as the reference standard. For each image, the absolute deviation of each algorithm’s estimate from the human reference was calculated, and deviations across all images were pooled to compute the mean absolute error (MAE).
In addition, two image-level performance metrics were evaluated: (i) the number of images in which one algorithm’s estimate deviated less from the human reference than the other (“Closer to Human”), and (ii) the number of images in which an algorithm’s deviation from the human reference was smaller than the within-human standard error (“Within Human SE”). Differences in counts between BlueNuclei and StarDist were assessed using binomial tests, reflecting paired binary outcomes for each image indicating which algorithm produced an estimate closer to the human reference
